# Nonequivalence of *Zfp423* premature termination codons in mice

**DOI:** 10.1101/2025.05.30.656936

**Authors:** Dorothy Concepcion, Catherine Liang, Daniel Kim, Bruce A. Hamilton

## Abstract

Genetic variants that introduce a premature termination codon (PTC) are often assumed equivalent and functionally null. Exceptions depend on the specific architectures of the affected mRNA and protein. Here we address phenotypic differences among early truncating variants of mouse *Zfp423*, whose phenotypes resemble Joubert Syndrome and Related Disorders (JSRD). We replicate quantitative differences previously seen between presumptive null PTC variants based on their position in the coding sequence. We show with reciprocal congenic strains that large phenotype differences between two PTC variants with the same predicted stop and reinitiation codons is due to the specific allele rather than different strain backgrounds, with no evidence for induced exon skipping. Differences in RNA structure, however, could influence translation rate across the affected exon. Using a reporter assay, we find differences in translational reinitiation between two deletion variants that corelate with predicted RNA structure rather than distance from the canonical initiation codon. These results confirm and extend earlier evidence for differences among *Zfp423* PTC variants, identify parameters for translational reinitiation after an early termination codon, and reinforce caution in the null interpretation of early PTC variants.

## INTRODUCTION

Exceptions to general rules can highlight new biology. We previously reported both an extensive allelic series of *Zfp423* mutations in mice and differences among inbred strain backgrounds in phenotypic outcomes of a single null mutation. On a co-isogenic FVB/NJ (FVB) inbred strain background, frameshift mutations with a premature termination codon (PTC) earlier in the open reading frame showed a trend toward less severe physical abnormalities (Deshpande *et al*. 2020), although the number of samples per variant was limited and potential mechanism was unclear. In parallel, a single presumptive null mutation that showed complete perinatal lethality on a C57BL/6J (B6) inbred background showed strain-dependent survival frequencies in backcross progeny from any of 13 other inbred strains (Alcaraz *et al*. 2020). More surprisingly, one early PTC mutation on the sensitized B6 background had detectable expression of a truncated protein and substantially less severe outcomes than adjacent PTC variants (including one resulting in the same frame-shifted stop codon) on the more tolerant FVB background. This contrast indicated that either relatively minor differences in these mutations, or differences in strain background with respect to RNA processing or translation, mediated the striking difference in phenotype that ran counter to what would be expected from equivalent mutations on those backgrounds. Here we test the replicability and causes of these unexpected differences and propose a mechanism.

*Zfp423* encodes a nuclear protein with an array of 30 C2H2 zinc fingers that contact both nucleic acids and an assortment of transcription factors, regulatory proteins, and DNA damage response proteins. It was first described in rat as an inhibitory binding partner to early B-cell factor (EBF) family proteins to block premature differentiation in progenitor cells (Tsai AND Reed 1997; Tsai AND Reed 1998; Cheng AND Reed 2007; Cheng *et al*. 2007; Roby *et al*. 2012). *Zfp423* is required in cerebellar granule cell progenitors for mitogenic response to SHH (Alcaraz *et al*. 2006; Hong AND Hamilton 2016). ZFP423 also functions as a differentiation factor in response to other intercellular signaling pathways, forming activating complexes with bone morphogenetic protein (BMP)-dependent SMAD proteins (Hata *et al*. 2000; Ku *et al*. 2006), retinoic acid receptors (Huang *et al*. 2009), and cleaved Notch intracellular domains (Masserdotti *et al*. 2010). ZFP423 has also been found in complexes with centrosomal protein CEP290 and poly(ADP-ribose) polymerase (PARP) in response to DNA damage (Ku *et al*. 2003; Chaki *et al*. 2012). Phylogenetic analysis showed strong constraint against amino acid substitutions at most residue positions across diverse vertebrate lineages and identified a possible 31^st^ zinc finger of C4 class (Hamilton 2020), subsequently supported by an AlphaFold2 structural model (Jumper *et al*. 2021; Tunyasuvunakool *et al*. 2021).

Mutation studies defined important roles for *Zfp423* in several developmental contexts. Consistent with the molecular perspective of ZFP423 participating in cellular differentiation pathways, loss-of-function mutations in mice produced malformations in midline brain structures, most prominently with agenesis or hypoplasia of the cerebellar vermis (Alcaraz *et al*. 2006; Warming *et al*. 2006; Cheng *et al*. 2007) and associated choroid plexus (Casoni *et al*. 2020), as well as defects in olfactory epithelium (Cheng AND Reed 2007), and in both developmental and regenerative adipogenesis (Gupta *et al*. 2010; Plikus *et al*. 2017). Human *ZNF423* expression also had prognostic value in human cancers across several studies, including neuroblastoma (Huang *et al*. 2009), leukemias (Miyazaki *et al*. 2009; Harder *et al*. 2013; Iglesias *et al*. 2021), breast cancer (Ingle *et al*. 2013; Brentnall *et al*. 2016; Cairns *et al*. 2017; Bond *et al*. 2018; Wang *et al*. 2019), glioma (Signaroldi *et al*. 2016), and stomach cancer (Nation *et al*. 2021). *ZNF423* mutations have also been implicated in ciliopathy disorders with cerebellar vermis hypoplasia and impaired DNA damage response (Chaki *et al*. 2012). Consistent with the idea that ZFP423 functions as an integrative node for multiple lineage- and signal-dependent pathways, both the anatomical phenotypes and null-allele viability show effect of genetic modifiers between different inbred strain backgrounds in mice. In particular, C57BL/6 backgrounds showed complete perinatal lethality for null mutations and more penetrant outcomes for a hypomorphic gene-trap allele in contrast with other strains (Alcaraz *et al*. 2011; Alcaraz *et al*. 2020). Phenotypes of different *Zfp423* mutations on a single background depend on domain composition and quantitative level of residual protein expression (Alcaraz *et al*. 2006; Casoni *et al*. 2017; Deshpande *et al*. 2020). In our previous work with 50 different *Zfp423* variants (Deshpande *et al*. 2020), the first PTC variant with respect to the open reading frame, D78Vfs*6, had residual protein expression and was substantially milder than all other PTC variants. However, slightly milder outcomes from adjacent PTC variants prompted reviewers and others to ask whether those PTC variants that lacked detectable protein expression (below ∼5% detection threshold) might nonetheless have residual function in contrast to PTC variants later in the open reading frame.

Here we revisited PTC mutations that had unexplained differences in phenotype with data from an additional set of animals. We found that subtly milder phenotypes of PTC variants early in the open reading frame compared with those later in the open reading persisted, despite protein levels below detection. We conclude that the overall effect is likely real, but small in magnitude. For the single PTC variant that had detectable protein expression, we found through reciprocal congenic strains that allele composition rather than unlinked variants in the genetic background was responsible for its sharp distinction from other PTC variants, including an adjacent variant with the same predicted stop and potential translational reinitiation codons. We found no evidence for induced exon skipping that could explain difference in protein level. Using a dual luciferase reporter assay we found that loss of a predicted stem loop structure increased translational reinitiation and explained most of the variance between D78Vfs*6 and other early PTC variants. We found that other early PTC variants had reporter expression above background, consistent with low protein expression from those alleles below the limit of detection for brain lysate Western blots.

## RESULTS

### Quantitative phenotype differences among PTC variants by position

Motivated by response to our earlier work (Deshpande *et al*. 2020), we tested whether differences between *Zfp423* PTC variants early (exon 3) vs. later (exon 4) in the open reading frame were reproducible, using the same morphological measures with new subjects at more advanced backcross generations and with a different investigator making the measurements. The major RNA isoform includes eight coding exons, with the majority of the open reading frame in a 3.2-kb exon 4 (Figure 1A). Variants used for this study are given in Table 1 (variant positions have been updated to reflect consensus RefSeq proteins, see Materials and Methods) with a focus on comparing the earliest frameshifts relative to protein functional domains (Figure 1B) with respect to anatomical phenotypes including body weight and simple neuroanatomical measures made from dorsal surface view and coronal forebrain or sagittal hindbrain block face views (Figure 1C). Excluding D78Vfs*6 as an outlier, other PTC variants in exon 3 (H96Wfs*4 and H96Vfs*29) and at the beginning of exon 4 (L133Rfs*23) had slightly milder presentation in weight and cortical area, with statistically non-significant or unreplicated trends in cerebellum midline sagittal area, mean cortical thickness, and maximum brain width (Figure 1 D, E). Only the anterior-posterior distance of cerebellum hemispheres had a nominal difference in the opposite direction, and only significant for one comparison across the replicate cohorts.

**Table 1.**
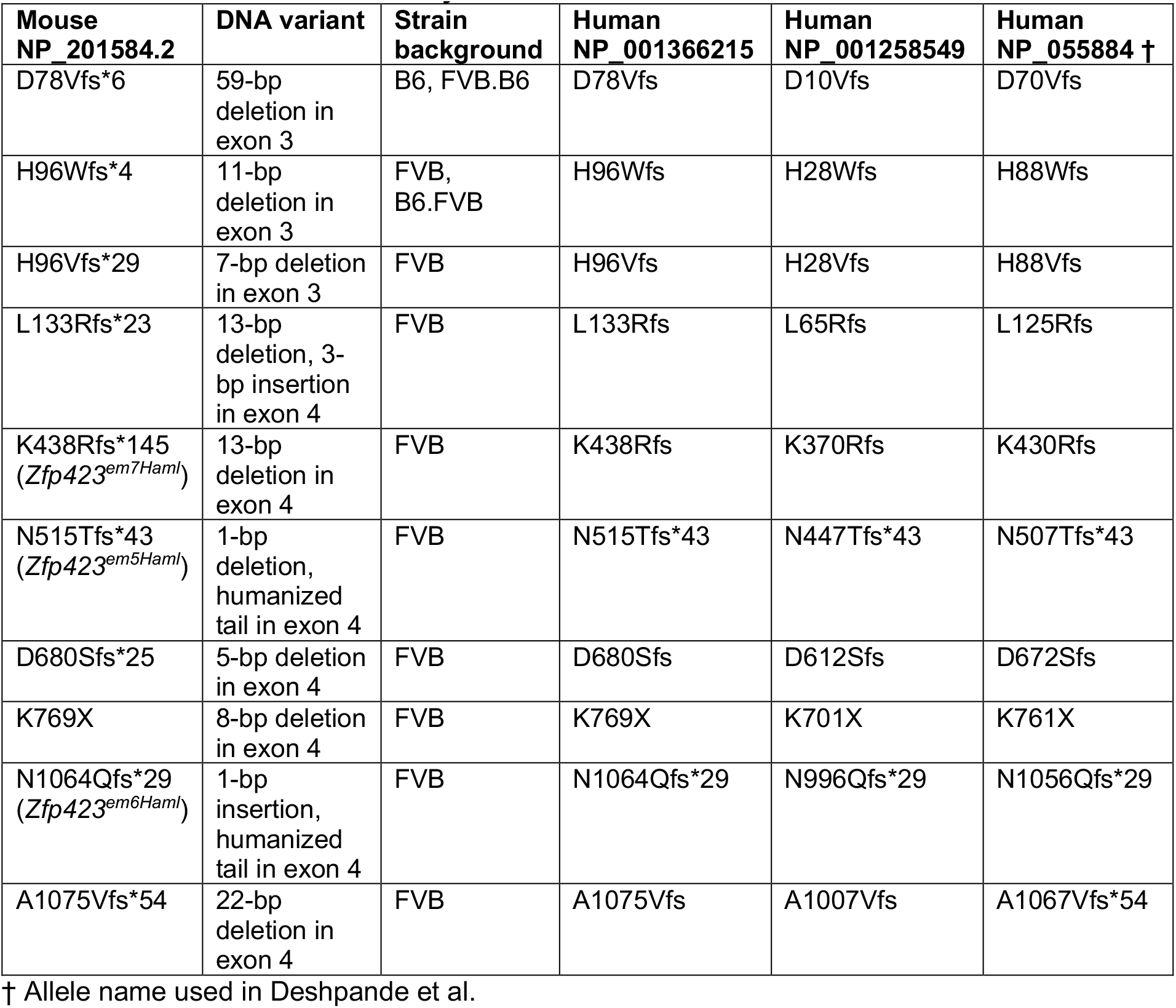
Variants used in this study.

**Figure 1.**
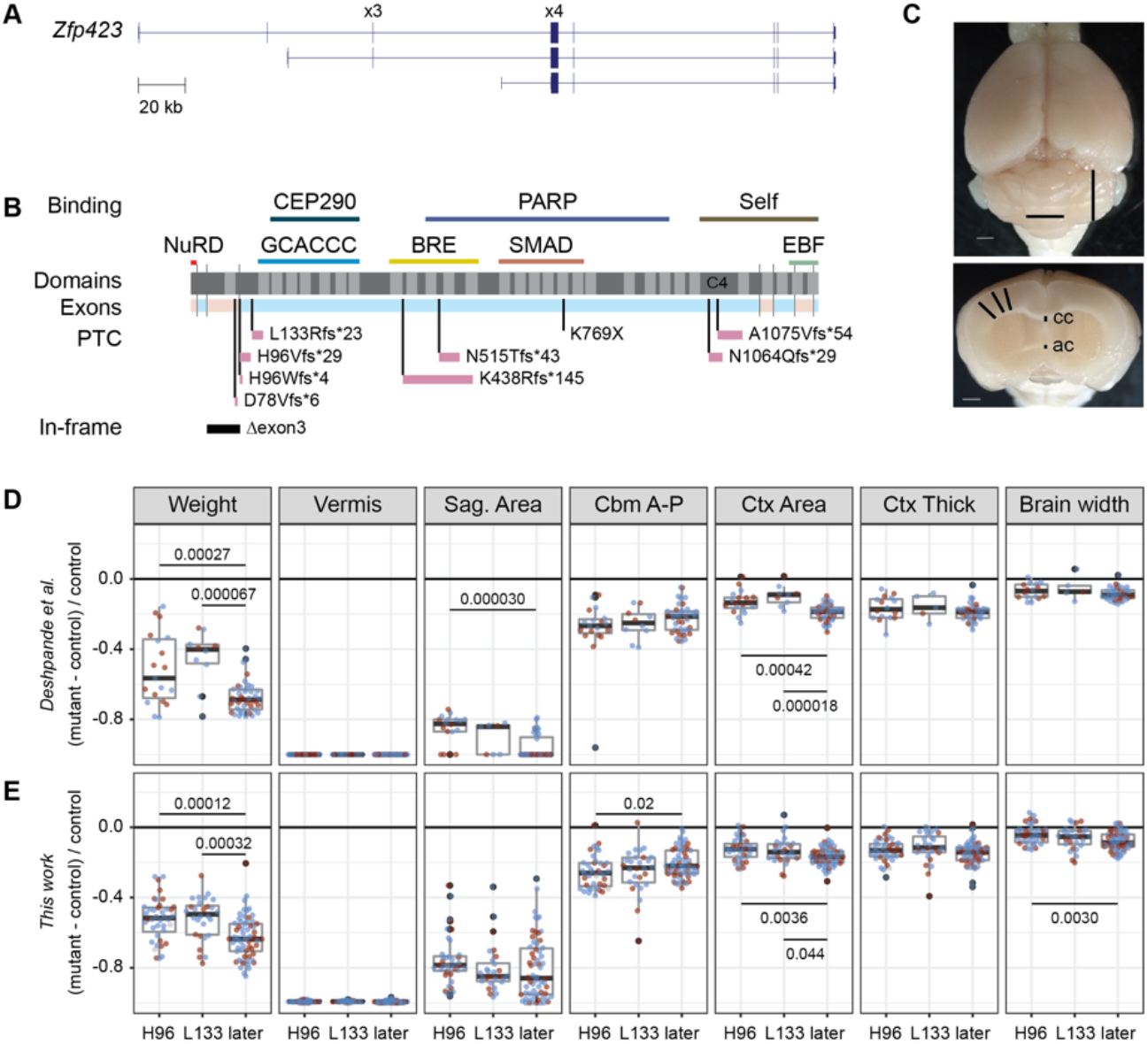
Unequal impacts of early vs. late premature termination codons among variants with protein levels below detection. (**A**) *Zfp423* gene structure showing constitutive exons 3 and 4. Adapted from UCSC genome browser. (**B**) Schematic of ZFP423 primary protein structure shows known binding domains (colored lines above) relative to C2H2 zinc fingers (lighter gray) above PTC variants, with relative lengths of shifted reading frames (pink bars), and exon 3 deletion allele. (**C**) Surface and coronal block face images with lines to illustrate the vermis width, cerebellum hemisphere anterior-posterior length (top); cortical thickness at 45°, 30°, and 15° and thickness of (cc) corpus callosum and (ac) anterior commissure (bottom). Adapted from (Deshpande *et al*. 2020). (**D**) Replotted data from Deshpande *et al*. showed less severe impacts of early compared with later PTC variants for body weight, cerebellum sagittal area at midline, and cortical area from dorsal view in the original data set. (**E**) Data from new paired samples replicate non-equivalent effects on weight and cortical area. Lines with numbers show comparisons with significant (p<0.05) differences between groups by Tukey’s honest significant differences post-hoc test after significant ANOVA for the three groups. Tests with p<0.0055 were considered to survive Bonferroni correction for the nine anatomical measures used.

Measures of midline thickness of corpus callosum and anterior commissure were noisier and statistically non-significant. This supports a functional difference between early and late PTC variants in *Zfp423*.

### Allele composition, not strain background, drives differences among early PTC variants

The largest difference among PTC variants was between D78Vfs*6, created and maintained on a C57BL/6 background, and all other PTC variants, created and maintained on FVB/NJ. D78Vfs*6 (59 bp), H96Wfs*4 (11 bp), and H96Vfs*29 (7 bp) are all small deletions within exon 3 that resulted in frameshifts after initial open reading frames encoding 9.3, 11, and 14 kD peptides, respectively, each ending in a premature UGA (opal) stop codon (Figure 2A). Because D78Vfs*6 and H96Wfs*4 sequences predicted the same premature termination codon and the same potential reinitiation codon, at M126, we tested whether the difference in outcomes could be explained by difference in genetic background.

**Figure 2.**
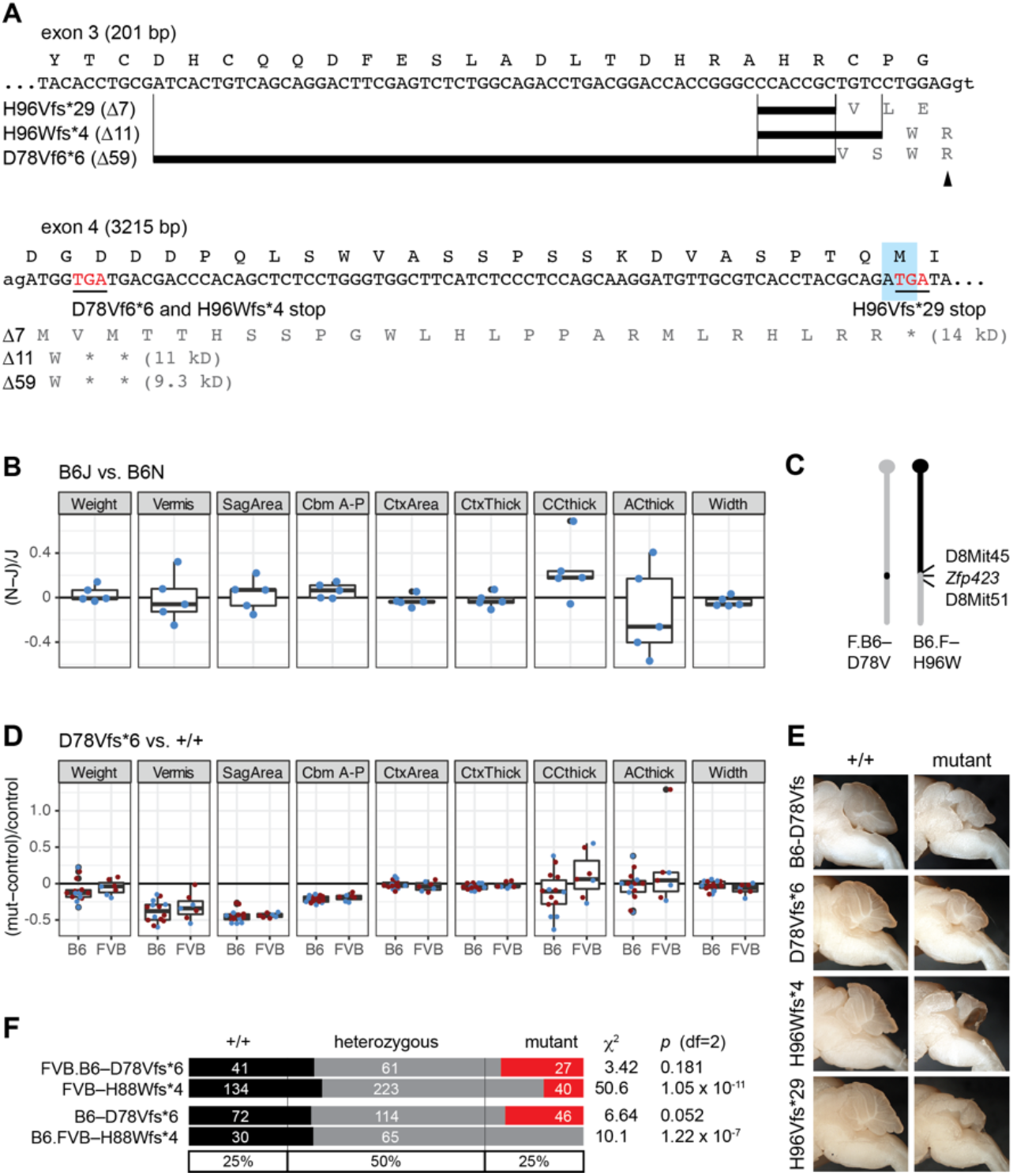
PTC variants early in the *Zfp423* open reading frame. (**A**) DNA sequences of three frame-shifting deletions near the 3’ end of exon 3. Single-letter amino acid translation shown above sense strand DNA sequence. Deleted bases indicated by thick black lines for each allele. Termination codons (underlined) and potential in-frame reinitiation codon (shaded box) within 5’ end of exon 4 are indicated, as are the predicted molecular weights of the potential 5’ translation products. Exon 3 splice donor and exon 4 splice acceptor sites are indicated with lower case. A frame-shifted arginine codon, AGA, that would be decoded by the n-Tr20_UCU_ tRNA (arrowhead) spans exons 3 and 4. (**B**) The C57BL/6J-specific modifier allele of n-Tr20_UCU_ does not have a major impact on D78Vfs*6 phenotype measures. (**C**) Extent of FVB (grey) and C57BL/6J (black) on chromosome 8 for reciprocal congenic strains. (**D**) FVB background does not make D78Vfs*6 more severe or comparable to H96Wfs*4 in brain measures. (**E**) Sample images from control littermate and exon 3 PTC variants on C57BL/6J (B6) and FVB/NJ congenic or coisogenic strain backgrounds. (**F**) H88Wfs*4 has lower survival frequency than D78Vfs*6 on either strain background, including complete lethality in C57BL/6J, comparable to other null alleles (Alcaraz *et al*. 2011; Alcaraz *et al*. 2020).

We found no sequence differences in the exons or intron-exon junction sequences between B6 and FVB for exons 3 and 4–neither in public data (Keane *et al*. 2011; Yalcin *et al*. 2011) nor by targeted resequencing during development of these alleles (Deshpande *et al*. 2020). Since D78Vfs*6 and H96Wfs*4 shared the same predicted stop codon after frameshift and the same potential reinitiation codon (M126, Figure 2A), this suggested that the effective difference must be either the length or content of the larger deletion or an effect of genetic background. Although *Zfp423* outcomes are generally more severe in C57BL/6 strains (Alcaraz *et al*. 2020), the C57BL/6J substrain used is known to harbor genetic modifiers of translation that might have allele-specific effects (Ishimura *et al*. 2014; Ishimura *et al*. 2016; Kapur *et al*. 2020).

C57BL/6J includes a variant neuronal-specific tRNA (*n-TRtct5*, also called *n-Tr20*_*UCU*_) that acts as a genetic modifier by stalling translation at AGA arginine codons and triggers an integrated stress response (Ishimura *et al*. 2014; Ishimura *et al*. 2016). This codon occurs both upstream of the D78Vfs*6 frameshift at position R62 and within the altered reading frame two codons before the terminator (mutant-specific R80 in D78Vfs*6; arrowhead in Figure 2A). To test whether this variant might suppress D78Vfs*6, we compared *Zfp423*-D78Vfs*6 phenotypes in a cross to a congenic strain that carried the wild-type B6N allele of *n-TRtct5*, comparing measures between sex-matched, D78Vfs*6-homozygous littermates that were homozygous for alternate alleles of *n-TRtct5*. We did not detect differences in any phenotype measure among five littermate pairs that would be consistent with the B6N *n-TRtct5* allele enhancing D78Vfs*6 phenotypes at a level that could explain the major allelic differences between PTC variants (Figure 1B).

To test the effect of strain background more broadly, we constructed reciprocal congenic strains to move the D78Vfs*6 allele onto FVB/NJ and the H96Wfs*4 allele onto C57BL/6J by marker-assisted breeding (Figure 2C). If a major component of the difference between B6–D78Vfs*6 and FVB–H96Wfs*4 derived from the B6 background suppressing effects of D78Vfs*6 (or FVB enhancing effects of H96Wfs*4), then the two alleles should be more similar when put on the same background. We observed the opposite. D78Vfs*6 phenotypes tended to be slightly milder in the FVB.B6 congenic line than on the original B6 background (Figure 2D). Incipient congenic B6.FVB-H96Wfs*4 failed to produce viable homozygotes at five or more generations of backcross (Figure 2E). Major anatomical differences between mutant alleles, including at the cerebellum midline, persisted among the exon 3 PTC variants on the FVB background (Figure 2F). These results, together, indicated that the milder phenotype of D78Vfs*6 relative to the other exon 3 PTC variants is attributable to differences in the allele composition rather than general or allele-specific modifiers in either genetic background.

### Exon 3 PTC variants do not induce significant exon skipping

If small deletions within exon 3 induced exon skipping, then the resulting mRNA would have a restored open reading frame, encoding a protein that lacked the first zinc finger. We previously showed that three distinct alleles that delete either the first zinc finger or all of exon 3 produced only very mild phenotypes (Deshpande *et al*. 2020). In contrast, translational reinitiation after either of the frame-shifted stop codons would produce a protein that lacked an aminoterminal peptide that recruits the repressive NuRD complex (Lauberth AND Rauchman 2006; Lauberth *et al*. 2007) as well as the first zinc finger. We assayed exon inclusion by reverse transcription-coupled polymerase chain reaction (RT-PCR) on cerebellum RNA using either of two primer sets that flank exon 3 (Figure 3A). Product lengths on gel electrophoresis showed that although we detected allele-specific exon 3 lengths, we did not see evidence for exon skipping in any of H96Vfs*29 (Δ7), or H96Wfs*4 (Δ11), or D78Vfs*6 (Δ59) compared with the exon 3-deletion allele (Δ345). To quantify the sensitivity of the assay, we used a serial dilution of either homozygous or heterozygous exon 3-deleted cDNA into control cDNA in the same PCR experiment and gel (Figure 3B). This showed a detection threshold of ∼0.5% of exon skipping. These results show that nonsense-mediated exon skipping cannot explain the ∼50% protein levels previously observed for D78fs*6 and, as the three small deletions largely overlap, are unlikely to explain residual function of the other exon 3 frame-shifting deletions.

**Figure 3.**
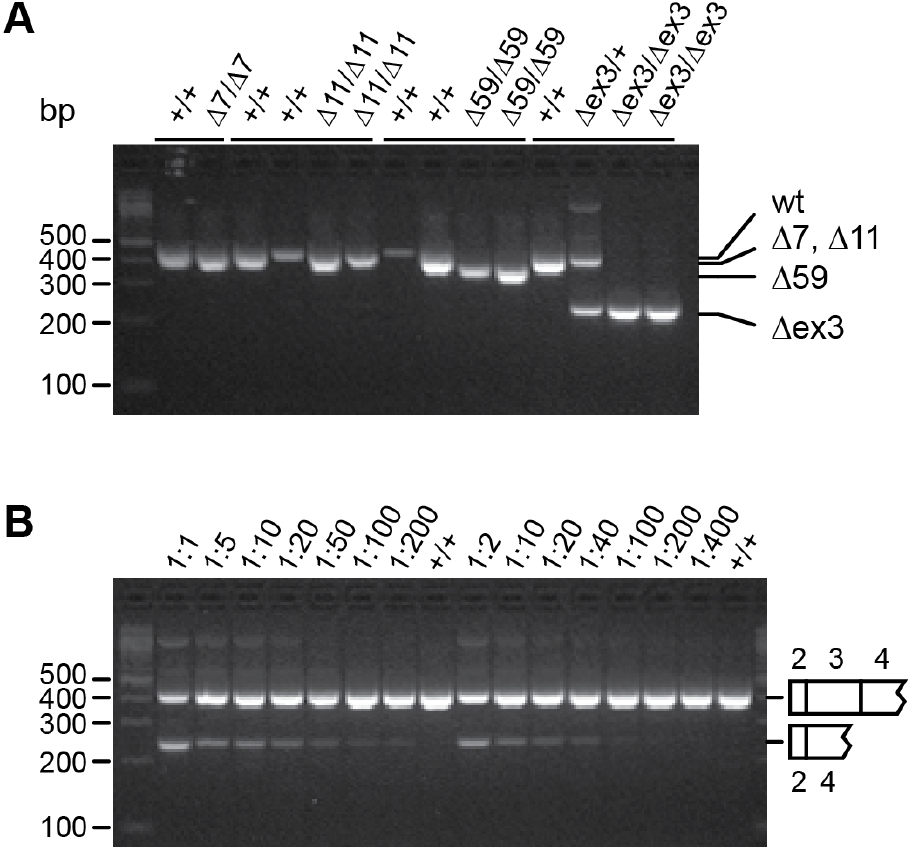
Lack of exon skipping around exon 3 PTC variants. (**A**) RT-PCR assay for exon 3 inclusion shows size variation due to deleted nucleotides but no evidence for exon skipping in comparison to an exon 3 deletion (Δ345). Similar results were obtained with two distinct primer sets. (**B**) Titration of Δ345 into a wild-type control shows RT-PCR detection sensitivity to ∼0.5% exon skipping. Assay run using the same master mix, PCR plate, and gel as (A). Left and right dilution series from different RNA samples.

### Exon 3 PTC variants differ in RNA secondary structure potential

If the difference between D78Vfs*6 and the other exon 3 variants is not attributable to strain background or exon skipping, then the most likely explanation would be a difference in translational reinitiation after the frame-shifted stop codon. Known factors that influence the likelihood of reinitiation include distance between first stop codon and the reinitiation codon, identity of the penultimate codon before the stop, distance from the upstream start codon to the reinitiation codon, and secondary structure of the mRNA (Kozak 2001; Bohlen *et al*. 2020; Dever *et al*. 2023). D78Vfs*6 and H96Wfs*4 have identical penultimate codons, stop codons, and potential reinitiation codons (Figure 1A), leaving distance from the upstream initiation codon and secondary structure (or a combination of the two) as possible factors. D78Vfs*6 placed the presumptive reinitiation codon 319 nucleotides from the canonical AUG, after an 81-codon upstream open reading frame (uORF), while H96Wfs*6 placed it slightly further at 367 nucleotides from the canonical AUG, after a 97-codon uORF–a 48 nucleotide or 15% increase in distance. The 59-bp deletion in D78Vfs*6 also removed most of an imperfect inverted repeat with strong potential for stem loop formation in the RNA (Figure 4A). RNAfold (Gruber *et al*. 2008; Lorenz *et al*. 2011) predicted a stem loop from this sequence with a -30.5 kcal/mol Gibbs free energy change (ΔG) for the optimal conformation, which is identical between mouse and human. By comparison, the 7-bp and 11-bp deletions in H96Vfs*29 and H96Wfs*6 retained predicted stem loop structures (Figure 4B), although with reduced predicted free energy change relative to the wild-type sequence (Figure 4C). This suggests that secondary structure might play a primary role in differentiating D78Vfs*6 from the other exon 3 PTC variants.

**Figure 4.**
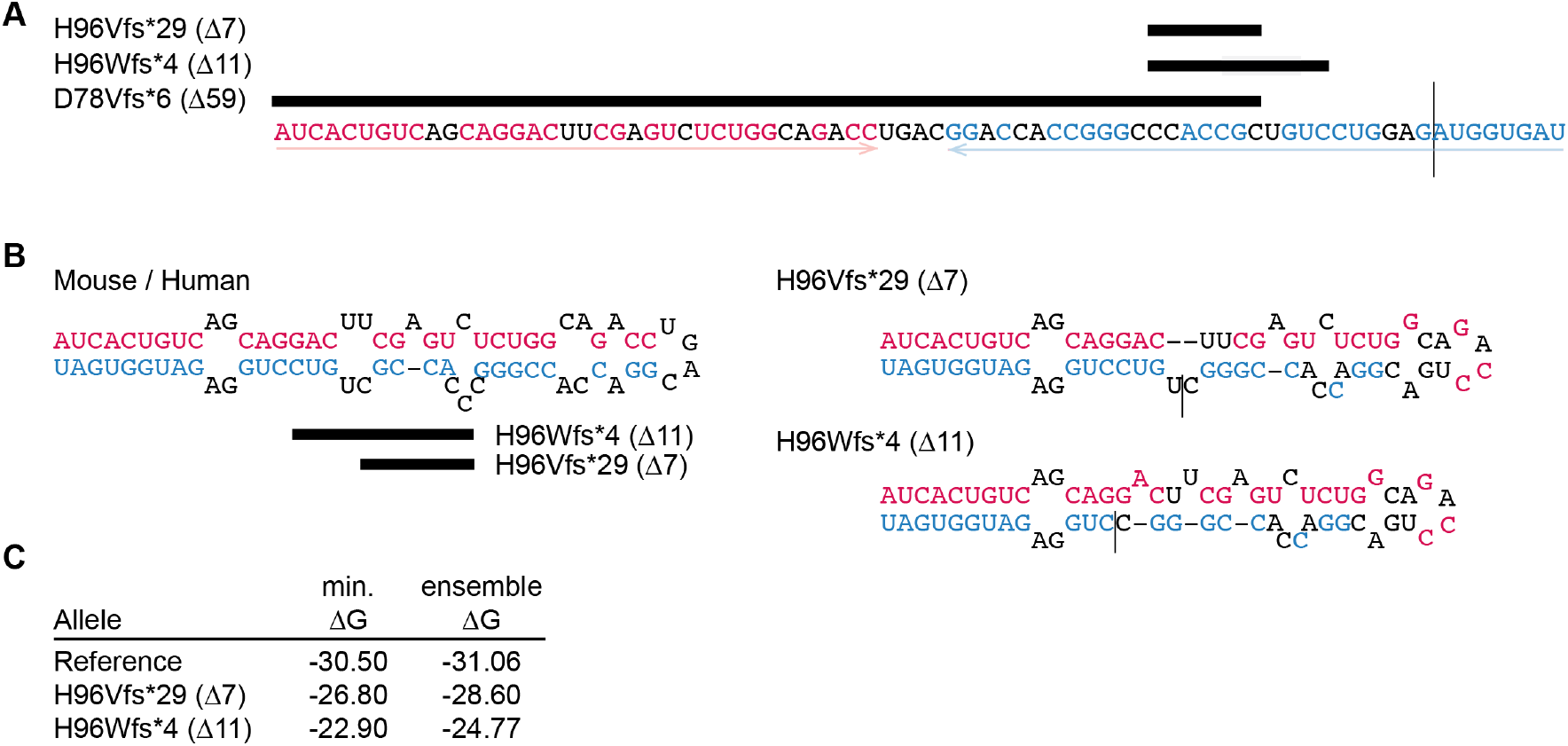
RNA stem loop before the frame-shifted stop codon differentiates exon 3 alleles. (**A**) Exon 3 deletion alleles relative to RNA sequence with a predicted stem loop. Vertical line, exon 3-exon 4 junction. Arrows indicate imperfect inverted repeat, with predicted paired bases shown in red (left arm) and cyan (right arm). (**B**) Predicted optimal structures of the mouse and identical human reference allele, H96Vfs*29 (Δ7), and H96Wfs*4 (Δ11), retaining the color coding from (A). (**C**) Predicted free energy (ΔG) for the optimal conformer (min.) and ensemble of conformer structures for each allele.

### Stem loop sequence differentiates expression levels of exon 3 PTC variants

To identify determinants of expression differences among the exon 3 frameshift variants, we developed a single-plasmid dual luciferase reporter assay (Figure 5A). An EF1α promoter drove expression of the *Zfp423* 5’ untranslated region (UTR) and open reading frame through codon 128 fused through a glycine/serine-rich linker to an in-frame Nano luciferase reporter, followed by the *Gapdh* 3’ UTR and polyadenylation signal with an additional synthetic polyadenylation signal before a *Pgk1* promoter driving firefly luciferase as a control reporter. In addition to a wild-type reference sequence and the D78Vfs*6, H96Wfs*4, and H96Vfs*29 variants, we synthesized novel variants to distinguish effects of nucleotide length between the canonical start and potential reinitiation codon from effects of disrupting the predicted stem loop (Figure 5B).

**Figure 5.**
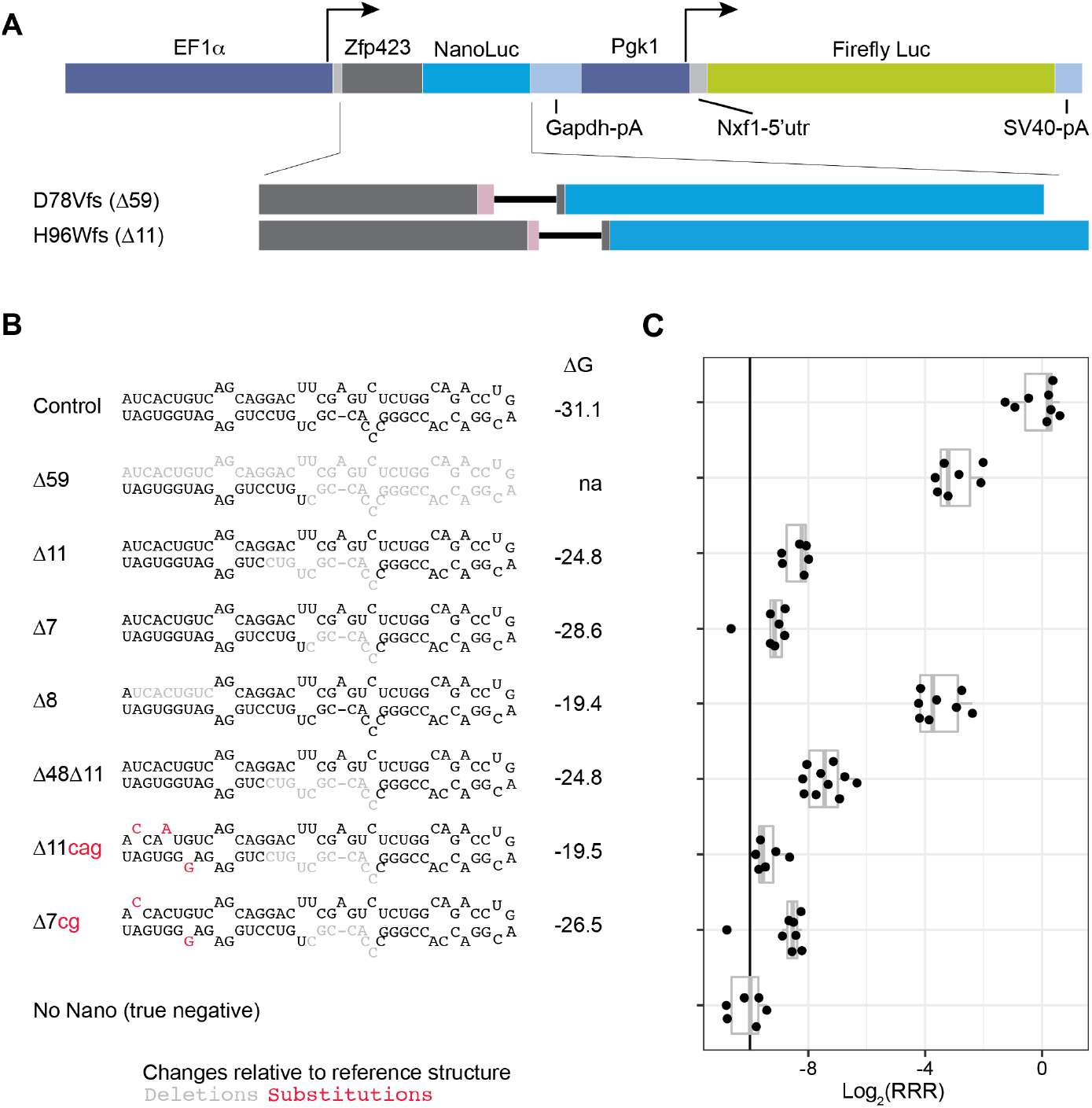
Luciferase reporter assay. (**A**) Dual luciferase reporter construct. EF1α promoter drives expression of *Zfp423* 5’ UTR and coding sequence through codon G128, fused to a Nano luciferase coding sequence (NanoLuc) followed by the *Gapdh* 3’ end with an additional polyadenylation signal. A *Pgk1* promoter drives expression of an *Nxf1* 5’ UTR and Firefly luciferase coding sequence (Firefly Luc) followed by an SV40-T 3’UTR and polyadenylation signal. Schematics below illustrate the D78Vfs*6 (Δ59) and H96Vfs*4 (Δ11) variants, show the short altered reading frame (pink) and distance (line) between shifted stop codon and the next in-frame AUG at M126. (**B**) Schematic illustration of tested variants relative to the predicted stem loop in reference sequence. Right, computed ensemble ΔG (RNAfold) for just the sequence shown, excluding deletions. Deleted positions in light gray, base substitutions in red. (**C**) Relative response ratio (see methods) for the Nano luciferase reporter relative to the Firefly luciferase for each construct. Each dot represents a separate transfection.

Each *Zfp423* variant along with the wild-type sequence as a positive control and negative controls that lack intact open reading frames for one or both luciferases were assayed on a single 96-well plate that included replicate DNA preparations for each construct (Figure 5C).

The dual-reporter assay recapitulates the order of expression levels observed in Western blots from wild-type and previously reported variants, distinguishes effects of deletion length from sequence composition, and supports low but non-zero expression from H96Wfs*4 and H96Vfs*29 variants. We detected strong but hypomorphic expression from the D78Vfs*6 (Δ59) allele reporter relative to the wild-type control, consistent with previous Western blot data from brain lysates (Deshpande *et al*. 2020), and much lower expression from H96Wfs*4 (Δ11) and H96Vfs*29 (Δ7) reporters. Importantly, a synthetic 8-bp deletion (Δ8), with a longer distance between AUG codons but deleting the perfect-match segment of the predicted stem loop had expression indistinguishable from Δ59. In contrast, deleting an extra 48 bp of sequence from a region upstream of Δ11 to create the same distance between AUG codons as Δ59 but with the same stem loop sequence as Δ11 had expression comparable to Δ11, as did nucleotide substitution variants with modest predicted effects on free energy (ΔG) of the stem loop. In an ANOVA model for all variants considered (p = 1.68 × 10^−8^), the D78Vfs*6 (Δ59) and 8-nucleotide deletion (Δ8) significantly differed from all other variants (p ≤0.05, Tukey HSD post-hoc tests) but not from each other (p>0.99). The H96Wfs*4 (Δ11) reporter was significantly above background level of the negative control (p = 6.4 × 10^−4^), while the H96Vfs*29 (Δ7) reporter was not, due to an outlier. These data support stem loop secondary structure as a primary determinant of reinitiation efficiency among these variants and support low level ZFP423 expression from the H96Wfs*4 and H96Vfs*29 alleles as a plausible mechanism for differentiation of *Zfp423* early PTC variants from those later in the open reading frame.

## DISCUSSION

Here we investigated unexplained phenotype differences among premature termination codon (PTC) variants caused by small, frame-shifting deletions in *Zfp423*. Our earlier work showed that D78Vfs*6 (a 59-bp deletion in exon 3) was an outlier both in phenotype and in producing a detectable truncated ZFP423 protein. We had expected that all other PTC variants would be equivalent null alleles, with the small differences among them attributable to noise and chance. To our surprise, the data showed a small but reproducible effect of position in which the next three PTC variants (H96Wfs*4 11-bp deletion, H96V*29 7-bp deletion in exon 3; L133Rfs*23 10-bp deletion near the beginning of exon 4) had reproducibly less severe weight and cortical area deficits relative to littermate controls, compared with other PTC variants further into exon 4 (Figure 1D, E). As this effect included a PTC at the 5’ end of exon 4 (3215 bp), but not PTC variants further in the exon, this seems unlikely to be explained by alternative transcripts (Figure 1A) or exon 3 skipping and may instead suggest a small amount of translational reinitiation with resulting steady-state protein level below our detection threshold. We cannot rule out a recurrent statistical anomaly or latent variable between allelic strains, although we took care to breed each line concurrently on the same vivarium rack, with boxes interspersed by allele, to minimize any environmental factor.

We had expected the much larger difference between D78Vfs*6 and other PTC variants to be explained by either genetic background (such as translational pausing related to mutation of a tRNA specific to a codon in the shifted reading frame ahead of the potential reinitiation AUG) or to frequent exon-skipping (exon 3 deletion preserves reading frame and had a milder presentation than even D78Vfs*6 in previous data). Data from crosses to congenic tRNA variant alleles argued against the specific tRNA hypothesis (Figure 2B). Data from reciprocal congenic strains showed that while strain background did affect severity across alleles, consistent with previous results (Alcaraz *et al*. 2011), it was not a significant component of the difference between exon 3 PTC variants (Figure 2D). A calibrated exon inclusion assay (Figure 3) further showed that exon skipping could not account for more than 0.5% of steady-state *Zfp423* transcript–too little to account for D78Vfs*6-derived protein levels, which we previously measured at ∼50% of littermate controls (Supplemental Figure 1 of (Deshpande *et al*. 2020)).

We infer from these data that D78Vfs*6 allowed reinitiation of translation after the premature termination codon, likely beginning at the next in-frame AUG codon, M126. Surprisingly, this was a sharp distinction from H96Wfs*6, which should have the same frame-shifted termination codon and the same reinitiation codon at the same distance from that stop. Both distance from the upstream start and mRNA secondary structure can influence reinitiation efficiency in a context-sensitive manner through time-dependent release of translation initiation factors from ribosomes, with longer translocation time before reinitiation reducing the probability of reinitiation (Kozak 2001; Bohlen *et al*. 2020; Dever *et al*. 2023). It is possible that reinitiation efficiency drops sharply between *Zfp423* exon 3 PTC variants purely as a function of distance (difference of 48 nucleotides, 15% of the distance between the canonical and likely reinitiation AUG codons). We noted, however, that *Zfp423* exon 3 and the first several bases of exon 4 included an imperfect inverted repeat with potential to form a 27-bp stem loop, of which the left arm is completely removed by D78Vfs*6, while only 7 paired bases from the right arm are removed by H96Wfs*6 (Figure 4). Secondary structures are known to slow the rate of translational elongation in mammalian cells (Kozak 2001) and the predicted stem loop sequence is identical between mouse and human and conserved across other mammals, as the ZNF423 amino acid sequence is highly constrained across all vertebrates (Hamilton 2020) and the amino acids encoded by the stem loop region have limited codon degeneracy. This is consistent with a recent finding that insertion of an exogenous inverted repeat into the 5’ untranslated region of *Zic3* suppressed translation from its canonical start codon (Bellchambers *et al*. 2025), but here applied to an endogenous sequence in the context of reinitiation after a premature termination codon.

Data from a reporter assay indicate that secondary structure rather than change in length between AUG codons is a primary determinant of expression differences among the early PTC variants and suggest residual reinitiation from low-expression exon 3 PTC variants might explain the modest phenotype difference we noted in mice. The dual-luciferase reporter we developed was highly sensitive and recapitulated the order, though not absolute values, of differences we previously reported for steady-state ZFP423 level in brain lysates from the corresponding mutant mice. The low-level reporter expression we found could not be attributed to spillover from the control luciferase or other nonspecific effects by comparison to a negative control plasmid that had a frameshift in the nano luciferase open reading frame. This resolved the paradoxical differences we previously saw among *Zfp423* PTC variants in mice and the modular design of the assay vector provides a well-controlled platform for testing reinitiation potential of other premature termination codon variants relative to matched controls.

Our results add to a growing recognition of non-null effects of early PTC variants. Pathogenic nonsense variants in the first exon of *PHOX2B* (codons 5-14) caused congenital central hypoventilation syndrome, but without the more severe presentations associated with later PTC variants or polyalanine expansion alleles, and with expression of an N-terminally truncated protein in cell models (Cain *et al*. 2017). In another cell culture transfection model, early PTC variants of *TP53* induced detectable reinitiation, with efficiency decreasing across length, for PTC variants up to 126 codons from the canonical AUG (Cohen *et al*. 2019); this is comparable to the subtle phenotypic improvements we found for *Zfp423* H96Wfs*6, H96Vfs*29, and L133Rfs*23 relative to later PTC variants *in situ*. At similar distance, a *KCNH2* PTC variant (T152Pfs*14) supported reinitiation at a noncanonical CUG codon, producing an N-terminally truncated protein with dominant negative properties (Park *et al*. 2024). Similarly, early PTC variants in *INPP4A* were recently reported with less severe neurodevelopmental consequences than later PTC variants, although difference in protein expression were not directly demonstrated (Rawlins *et al*. 2025). Our results together with recent literature suggest that early PTC variants are often not fully null and in that in some clinical settings therapeutics that increase the efficiency of reinitiation after a stop codon might have benefit.

## MATERIALS AND METHODS

### Mice

*Zfp423* mutant strains were previously described (Deshpande *et al*. 2020). Mutations made by editing FVB/NJ zygotes were maintained as co-isogenic lines by backcross for a minimum of eight generations prior to studies described here. The D78Vfs*6 mutation (previously referred to as D70Vfs*6) was originally made in C57BL/6N zygotes and was then maintained as a substrain congenic after ten generations of backcross to C57BL/6J. FVB/NJ and C57BL/6J were obtained from the Jackson Laboratory or bred in house from recently imported stock. *n-Tr20* congenic mice were previously described (Ishimura *et al*. 2014) and the gift of Dr. Susan Ackerman. Mice were maintained in a specific pathogen free (SPF) facility on 12 h light, 12 h dark cycle in high-density racks with HEPA-filtered air and ad libitum access to water and food (LabDiet 5P06), as previously reported. Sex-matched mutant and control littermate pairs were used as the unit of replication (n) for all experiments.

The PTC variants were originally described using residue positions relative to a human reference protein (NP_055884) that lacked the amino-terminal MSRRKQAK sequence, which is highly conserved among orthologs (Hamilton 2020) and mediates recruitment of the NuRD corepressor complex (Lauberth AND Rauchman 2006; Lauberth *et al*. 2007). We rename variants here relative to full-length mouse (NP_201584.2) and human (NP_001366215.1) reference proteins (Table 1).

### Anatomical measures

Measures from dorsal whole-mount and coronal and sagittal block face images were made as previously described (Deshpande *et al*. 2020), but measured by a different investigator (DC) than our previous work. Briefly, animals were perfused with phosphate buffered saline followed by 4% paraformaldehyde and brains were removed into fresh 4% paraformaldehyde at 4°C for 12–24 h. Brains were mounted on a black background under a stereomicroscope using standardized zoom and distance settings with a ruler in frame to verify scale and photographed with a digital camera. Paired samples were processed and imaged together. Anatomical features were measured in ImageJ. Vermis width was measured at the middle of lobule VII. Cerebellar hemisphere anterior-posterior length was measured as a vertical line drawn from the dorsal-most point. Coronal and sagittal block face preparations were made using a standard mouse brain matrix (Zinc Instruments) with the sample aligned anteriorly. Coronal cuts through the striatum were made at the rostral end of the optic chiasm.

Sagittal cuts through hindbrain were made at the midline. For cortical thickness, lines at 15, 30, and 45 degrees from the dorsoventral axis were drawn using the ImageJ ROI Manager, with each line perpendicular to the brain surface and ending at the corpus callosum; the average of these three lengths was taken as mean cortical thickness. The thickness of the corpus callosum and anterior commissure were measured at the midline. Width of the brain was measured with a horizontal line across the coronal surface at its widest point. Cortical area was measured from the dorsal surface image and cerebellum midline area was measured from sagittal block face image as a manually defined region of interest in ImageJ.

### RNA analysis

Cerebella were dissected in ice-cold phosphate buffered saline, transferred to Trizol reagent (ThermoFisher), and homogenized with a Polytron with standardized speed settings across samples. First strand cDNA was synthesized with SuperScript III (Invitrogen). RT-PCR for exon 3 inclusion was tested using PCRx reagents (ThermoFisher) with two different primer sets that produced equivalent results: CGCGATCGGTGAAAGTTGAA with GGATGAAGGACTTGTCGCAG (5’ primer spanning exons 1-2, 3’ primer in exon 4) and CCTCGGACTTCTCGCTGG with AGGCGGATGAAGGACTTGTC (5’ primer in exon 2, 3’ primer in exon 4). Primers were purchased from IDT.

### Dual luciferase assay

A 3.5-kb fragment comprising the wild-type *Zfp423* 5’ untranslated region and coding sequence through the Gly128 codon, a Gly/Ser-rinh linker, Nano-luciferase coding sequence, *Gapdh* 3’ untranslated region with an additional synthetic polyadenylation sequence, human *PGK1* promoter, *Nxf1* 5’ untranslated region, and firefly luciferase coding sequence was commercially synthesized at Twist Biosciences and cloned into a pTwist-EF1a-Puro backbone linearized between the EF1a promoter and SV40 3’ end and polyadenylation signal with EcoRI and NheI, using HiFi assembly (New England Biolabs). Both luciferase coding sequences were modified to remove unwanted restriction sites to allow modular replacement of any functional component as needed. A single base clone was selected based on whole-plasmid sequence validation (Plasmidsaurus). Additional clones that failed sequence validation due to indels in either nano luciferase or firefly luciferase open reading frames and a single clone with both luciferase coding sequences deleted were used as specificity controls. Synthesized variant *Zfp423* fragments replaced wild-type sequence in the base clone after digestion with EcoRI and BamHI and HiFi assembly of synthetic fragments. All plasmids used were verified by whole-plasmid sequencing.

For each construct, plasmid DNA was prepared from at least three replicate cultures using Zyppy Plasmid Miniprep kits (Zymo Research Corporation). Plasmids were transfected into P19 cells in a 96-well dish using Lipofectamine 2000 and processed 18 hours after transfection. Assays were conducted using Dual-Luciferase reagents (Promega) according to manufacturer’s protocol and light emission measured on a Tecan Spark (luminescence mode, 360-700 nm wavelength, 500 ms). Raw measures were adjusted by subtracting the minimum values of negative control plasmids for each reporter. Relative Response Ratio (RRR) was calculated by using the ratios of adjusted nano and firefly luciferase values; RRR = (sample ratio - negative control ratio)/(positive control ratio - negative control ratio), using the minimum ratio among replicates that express only firefly luciferase as the negative control and the mean of wild-type replicates as the positive control. Log2 transformed values of RRR were used for visualization and ANOVA models.

### Statistical analyses

All statistical tests were conducted in R 4.1.3 either alone or through RStudio. ANOVA (aov) models used measures as a function of variant and pairwise comparisons between groups were assessed with Tukey’s honest significant difference (TukeyHSD) test.

## ACKNOWLEDGEMENTS

We thank Dr. Susan Ackerman for *n-Tr20* congenic mice and helpful conversations. We thank Ethan Satoda for assistance with genotyping. This work was supported by grant R01 NS097534 from the National Institute of Neurological Disorders and Stroke.

